# Cancer Cell Adaptation to Cycling Hypoxia: Integrating Mathematical Modeling with Experimental Validation of HIF-1 Dynamics

**DOI:** 10.1101/2025.04.06.647478

**Authors:** Xihua Qiu, Yamin Liu, Paola Vera-Nicola, Eran Agmon, Kshitiz, Yasir Suhail

## Abstract

The adaptive response of cancer cells to hypoxia, a key microenvironmental factor in solid tumors, is intricately orchestrated by Hypoxia-inducible factor 1 (HIF1). Evidence is accumulating on oxygen dynamics in tumor cores, showing that hypoxia is frequently unstable, or cycling, associated with specific phenotypic outcomes for the cancer. Transcriptomic analysis show that most gene expression changes in cycling hypoxia lie in between the change caused by stable hypoxia, suggesting multi-cycle averaging of dosage in the oxygen tension, and likely HIF-1 induced transcription. However, small subset of genes show an oscillation/cycling hypoxia specific response, suggesting that the transcriptional machinery of these genes may interpret cycling HIF-1 activity differently from stably high HIF-1 activity. Here, we model a gene regulatory circuit, the incoherent feed-forward loops (IFFLs) to explore parameter regimes where oscillatory specific transcription is plausible. In these IFFL models, HIF-1 regulates gene transcription of a target gene directly, as well indirectly via another transcription factor with an opposite effect on gene transcription. We have identified several plausible parameter regimes where oscillation specific gene expression can occur, with potentially significant effect on cancer growth and progression. Finally, we experimentally confirmed that HIF-1 can form IFFLs with two key transcription factors p53, and Notch1, resulting in cycling hypoxia specific gene expression linked to breast cancer progression and poor prognosis. Our models mechanistically reveal how temporal fluctuations in the tumor microenvironment may directly inform downstream transcription, identify hitherto unknown HIF-1 driven mechanism of cancer progression contributing to emergent tumor heterogeneity.

## Introduction

As tumors grow, they frequently outstrip their supply of oxygen, leading to development of hypoxic regions. Hypoxia is a key microenvironmental factor in solid cancers, influencing many aspects of cancer growth, metastasis, and drug resistance^1,2^. The cellular response to hypoxia is regulated by HIF-1 (Hypoxia-inducible factor 1), a transcription factor which acts as the master regulator of cellular responses to low oxygen levels (hypoxia)^3,4^. HIF-1 is a heterodimer, composed of the oxygen tension regulated HIF-1-alpha, and constitutively expressed HIF-1-beta, which is stabilized in low oxygen conditions, and acts as a key transcription factor (TF), regulating the expression of numerous genes involved in angiogenesis, metabolism, and cell survival to adapt to oxygen deprivation^5^. Consequently, the advent of hypoxia can have a profound effect on nearly all steps in the metastatic cascade^6^.

In the recent decade, development of *in vivo* imaging technologies have revealed that hypoxia is frequently not temporally stable, leading to fluctuations or cycling of oxygen tension^7-9^. Our own works, as well as others, have shown that cycling, or oscillatory, hypoxia can result in gene expression patterns associated with low patient survival, altered tumor inflammation, and stromal activation^7,10-13^. Furthermore, we had shown that stability of HIF-1 itself could fluctuate due to accumulated lactate in the tumor milieu, which facilitates non-canonical degradation of HIF-1 via chaperone mediated autophagy^14,15^. Crucially, while HIF-1 is known to suppress cell proliferation both transcriptionally by elevating tumor suppressor p21 and p27^16^, or counteracting the oncogenic effect of Myc^17^, as well as non-transcriptionally^18^, oscillatory/cycling hypoxia allowed the cells to continue to proliferate^10,14^, partially addressing the conundrum why tumors continue to proliferate in hypoxia.

Most of the effects of cycling hypoxia previously reported, as well as our own reported findings, indicate that the downstream HIF-1 driven gene expression is averaged for cycling hypoxia in between normoxia and stable hypoxia. A plausible explanation is that cycling hypoxia elicits a “sub-hypoxic” transcriptional response averaged over multiple cycles. This sub-hypoxic response, wherein the gene expression in cycling hypoxia is in between that of normoxia and stable hypoxia explains many of the observed phenomena, including increased cell proliferation, respiration etc.

However, gene expression analysis of HeLa^14^, MDA-MB-231^10^, as well as single cell RNAseq analysis of a population of various cell types in breast cancer tumor microenvironment showed many genes that had cycling/oscillatory hypoxia specific gene expression^19^. These genes unexpectedly show either a directionally congruent but more extreme response to stable hypoxia, or even opposite to stable hypoxia. Many of these genes could have a profound effect on cancer progression, and considering that they are induced specifically by cycling hypoxia, may provide hitherto unknown HIF-1 driven mechanisms of cancer progression.

How could a gene expression be driven specifically by cycling hypoxia, more than, or oppositely to stable hypoxia? Here, we model these patterns using Incoherent Feed Forward Loop (IFFL) motifs in the gene transcriptional machinery involving HIF-1 as the master transcriptional regulator^20^. IFFLs have been previously shown to be a strongly selected motif in prokaryotic transcriptional network^20^, and can differentiate variable signal from stable signal^21^. As HIF-1 is a master TF, regulating the expression of many other TFs with their own downstream gene expression effect, IFFLs could easily form where HIF-1 and HIF-1 induced gene product TF together regulate a target gene.

We analytically modeled various IFFL types, which could be formed between HIF-1, and a target gene with incorporation of another HIF-1 driven TF. Mathematical modeling revealed plausible regimes where cycling/oscillatory HIF-1 specific gene expression patterns could arise. Surprisingly, we found that a single IFFL subtype could give rise to many different cycling/oscillatory hypoxia specific gene expression patterns we identified in gene expression data in triple negative breast cancer MDA-MB-231 cells.

Finally, we found two key regulators, which could potentially act as candidate TFs to create IFFLs with HIF-1, the tumor suppressor p53, and Notch1, regulating expression of several oscillation specific genes. Measuring gene silencing for TP53 and NOTCH1, and measuring RNA levels by RNAseq in normoxia, stable hypoxia, and cycling hypoxia in MDA-MB-231 cells, we found many genes which form an IFFL gene regulatory circuit with HIF-1 along with either p53, or Notch1. Our mathematical model not only provides a mechanistic understanding of how gene expression patterns could emerge specifically in response to cycling hypoxia, but also demonstrate that microenvironmental dynamics can translate into specific transcriptomic, and therefore phenotypic responses in cancers.

## Results

### Gene expression in triple Negative Breast Cancer cells exhibit cycling hypoxia specific patterns

To ascertain the extent of gene expression changes in response to stably high (H) and cycling hypoxia (O) vs normoxia (N) in triple negative breast cancer cells (TNBCs), we used MDA-MB-231 cells and treated them with normoxia, hypoxia, and cycling hypoxia for 48 hours (**Figure 1A**). TNBCs frequently develop hypoxic cores, with peripheral regions with suboptimal perfusion experiencing fluctuating or cycling hypoxia (**Figure 1A**)^7,9^. As molecular oxygen has extremely low solubility in aqueous solutions^22^, dynamic oxygen stimulation is very difficult to control precisely, with soluble pO_2_ being highly sensitive to height of medium, air pressure, and many parameters. We therefore cultured MDA-MB-231 cells on gas permeable surfaces^10^, exposed from the bottom to the atmospheric oxygen, so that any change in the atmospheric pO_2_ is directly sensed by the cultured cells from the bottom of the plate.

**Figure 1.**
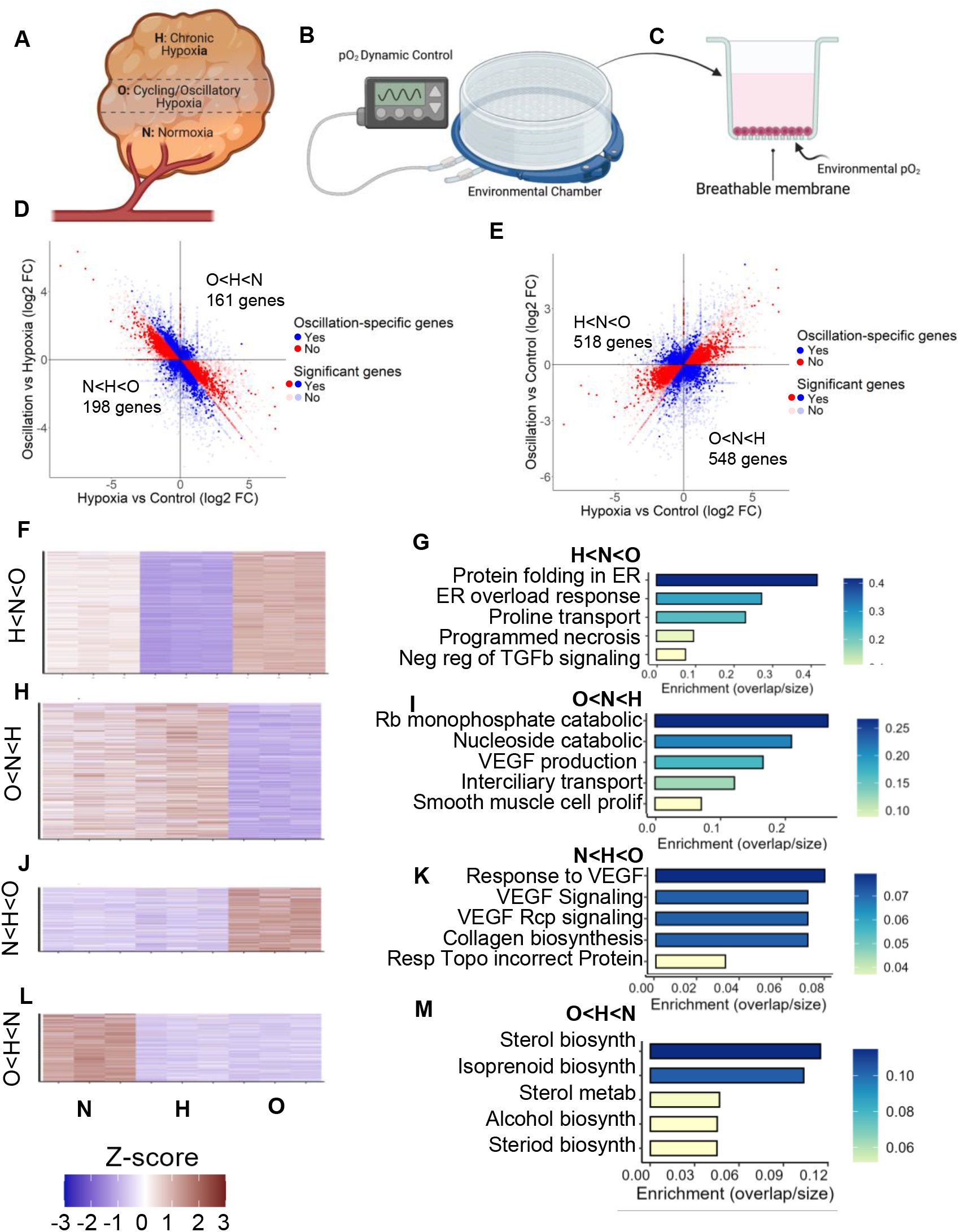
Cycling/Oscillatory hypoxia results in a specific pattern of gene expression in MDA-MB-231 distinct from stable/chronic hypoxia. **(A)** Schematic showing a sub-optimally vascularized solid tumor with regions of chronic hypoxia, cycling hypoxia, and normoxia in a tumor due to the spatial distribution of vasculature. (B -C) Experiment setup to control hypoxic dynamics in cell culture. (D) Scatter plot showing relative differential expression (in log2 fold change) between O vs H (y-axis), and H vs N (x-axis); Quadrants show gene expression following the oscillation specific pattern (H<N<O) top left, (O<N<H) bottom right. (E) Scatter plot showing relative differential expression (in log2 fold change) between O vs N (y-axis), and H vs N (x-axis); Quadrants show oscillatory specific patterns (N<H<O) bottom left, and (O<H<N), top right. (F-M) Heatmaps (left column) and top enriched pathways (right column), for genes with oscillation specific patterns H<N<O, O<N<H, N<H<O, and O<H<N respectively.

Cultured cells were kept in a sealed chamber connected to a dynamic gas mixing controller, and subjected to normoxia (N: atmospheric O2, ∼20%), hypoxia (H: 1% O_2_), and cycling/oscillatory hypoxia (O) in between N, and H with a period of 90min for a total of 48 hours (**Figure 1B-C**). Cells were immediately lysed, RNA collected, and sequenced. Differential gene expression analysis revealed that hypoxia expectedly had a dramatic effect on gene expression vs normoxia. With the adjusted p-values set for significance of 0.05 for all three fold changes, we counted the number of significant oscillation -specific genes in different quadrants (**Figure 1D-E**). As expected from our default hypothesis, for most genes, cycling hypoxia resulted in an averaged response in between normoxia, and stable hypoxia for most genes (**Figure 1D-E**). Surprisingly, we also found a large number of genes with cycling hypoxia specific changes in expression, not expected from the averaged response between normoxia and stable hypoxia (**Figure 1D-E**).

We categorized the patterns of cycling hypoxia specific gene expression in 4 broad categories defined by the significant differences between the three conditions arranged in terms of their tpm values. The first two patterns, O<H<N, and N<H<O consist of genes whose expression in cycling hypoxia was more extreme than in stable hypoxia (**Figure 1D**), while the other category included H<N<O, and O<N<H genes where cycling/oscillatory hypoxia had an opposite effect than stable hypoxia (**Figure 1E**). In none of the cases, the expression pattern of genes are explainable by the hypothesis that cycling hypoxia results in a sub-hypoxic an averaged response between normoxic and hypoxic states.

Gene set enrichment analysis of O genes revealed some phenotypic coherence. The 518 genes which showed a downregulation in stable hypoxia, but increased in cycling hypoxia (**Figure 1F**) were enriched in Protein folding response (**Figure 1G**), as we have previously reported, and shown to be linked to low patient survival^10^. Gene ontologies (GO) enriched in 548 genes which were decreased in cycling hypoxia, oppositely to stable hypoxia were of variable types, including ribosomal and nucleoside catabolic processes, as well as VEGF production (**Figure 1H-I**). Interestingly, among genes where cycling hypoxia showed a directionally similar, but an even more extreme response than stable hypoxia showed activation of GOs associated with angiogenesis, as well as collagen biosynthesis (**Figure 1J-K**). Cycling hypoxia resulted in decreased expression of genes associated with various biosynthetic pathways (**Figure L-M**), similar to stable hypoxia, although directionally more extreme. Overall, our data with MDA-MB-231 cells show a large number of genes which had a specific effect on expression by cycling/oscillatory hypoxia vs stable hypoxia, suggesting an indirect gene regulatory mechanism which parses variable signal differently from stable signal.

### Incoherent Feed Forward Loops (IFFL) driven by HIF-1 as plausible regulatory units for cycling hypoxia specific gene expression

We sought to understand how do so many genes specifically respond to cycling hypoxia, even oppositely to stable hypoxia. Considering that gene expression regulatory networks usually feature high interconnectedness and complexity, many genes are regulated through complex networks of transcription factors^23^. These networks include positive and negative feedback loops, among other sophisticated regulatory mechanisms^24^. Based on this information, we might speculate that a significant proportion of human genes—potentially several thousand up to over ten thousand—could be modeled using differential equations involving transcription factor regulation^25^. HIF-1 is a master transcriptional regulator, transcribing a large number of genes, many of which code for transcription factors^26^, creating a potential for formation of multi-protein regulatory circuits.

Among the many regulatory mechanisms, incoherent feedforward loops (IFFLs) have been mathematically described as being capable of discriminating between variable and stable signals^27,28^. Furthermore, IFFLs have been shown to be evolutionarily selected motifs in transcriptional regulatory networks in many species^28^. We there asked if IFFLs with HIF-1 being the driver TF may explain some of these patterns.

Feed forward loops (FFLs) are a general class of motifs composed of 3 elements, wherein a TF X that regulates a second TF, Y, and both X and Y regulate the target gene Z^20^. FFLs could be coherent, or incoherent, based respectively on the congruency, or incongruency of the parallel arm connecting the regulating TF X to the target Z (**Figure 2A-B**). IFFLs have been previously shown to be able to discriminate between variable and stable input signal X^27,29^. The IFFL consists of two parallel paths between X and Z, both being oppositely activating. The direct path consists of a single directed edge, and the indirect path is a cascade of two directed edges, such that their product of activating sign is opposite to the direct edge. It is to be noted that IFFLs unlike FFLs, are highly selected in evolution, and are considered as a gene regulatory motif^20,27^. There are 4 potential IFFLs (**Figure 2B**).

**Figure 2.**
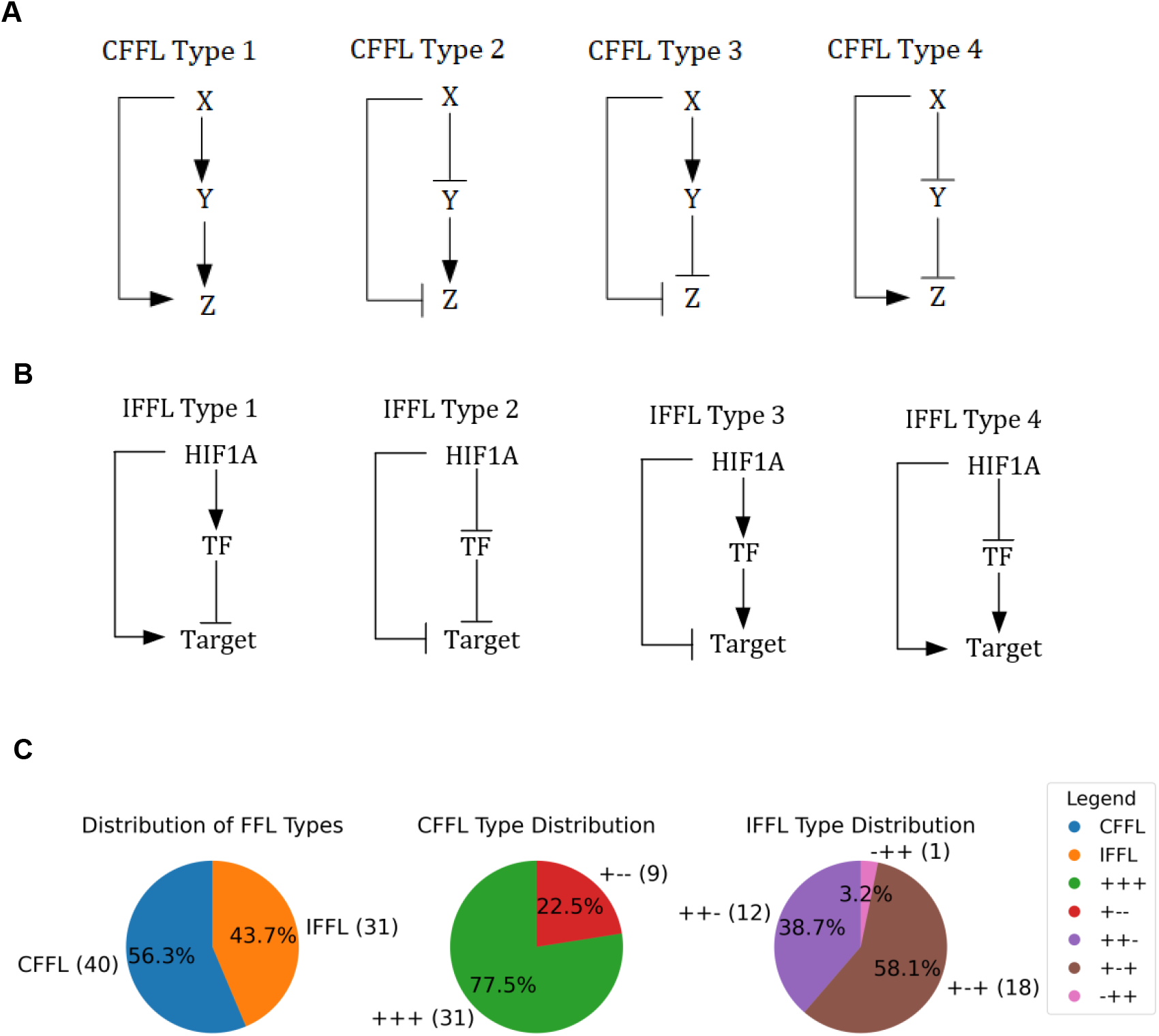
Gene regulatory feed forward loops involving activating and inhibitory interactions. (A) All possible coherent feed forward loops. The overall effect of driving gene X on the target gene Z is in the same direction from both the direct interaction (X -> Z) and the indirect interaction (X -> Y -> Z). (C) All possible incoherent feedforward loops, where the direct and indirect effects of X on Z are in opposite regulatory directions (activating and inhibitory). (C) Relative distribution of FFLs, both coherent (CFFLs) and incoherent (IFFLs) identified in MDA-MB-231 gene expression data obtained in the conditions: normoxia (N), stable hypoxia (H), and cycling/oscillatory hypoxia (O).

We analytically modeled each of them to test the possibility of obtaining cycling/oscillatory hypoxia (O) specific gene expression patterns reminiscent in the data collected in MDA-MB-231 cells in N, H, and O conditions. According to relevant literature, we have confirmed 31 IFFLs and 40 CFFLs that are regulated by HIF-1 (**Figure 2C**). Among them, the majority of the CFFLs (77.5%) belong to CFFL type 1 (+ + +), while the remaining 22.5% fall under CFFL type 3 (+ − −). Similarly, for the IFFLs, type 1 (+ − +) accounts for 58.1%, followed by type 3 (+ + −) at 38.7%, and a small proportion (3.2%) is classified as type 4 (− + +).

### Mathematical modeling of IFFLs

We mathematically modelled all four IFFLs types with ordinary differential equations. As an example, we discuss the IFFL type 3 gene circuit mediated driven by HIF-1 (**Figure 3**), a regulatory module where HIF-1 activates a secondary transcription factor (TF), which, in turn, regulates a downstream target gene. Simultaneously, HIF-1 directly downregulates the same target gene, creating a balance between activation and inhibition (**Figure 3A**). For nonlinear interactions at each regulatory edge, this circuit shows regimes wherein the target gene expression is driven selectively by cycling hypoxia. Equation (1) describes the expression of the intermediary transcription factor (TF) being driven by HIF-1 binding on its enhancer region, with the nominal production rate of *β*_*TF*_, and degradation rate of α_TF_ that is linear with the amount of TF present. Equation (2) describes our model with the target gene expression being driven by TF binding on its enhancer region and HIF-1 binding on its silencer region.

**Figure 3.**
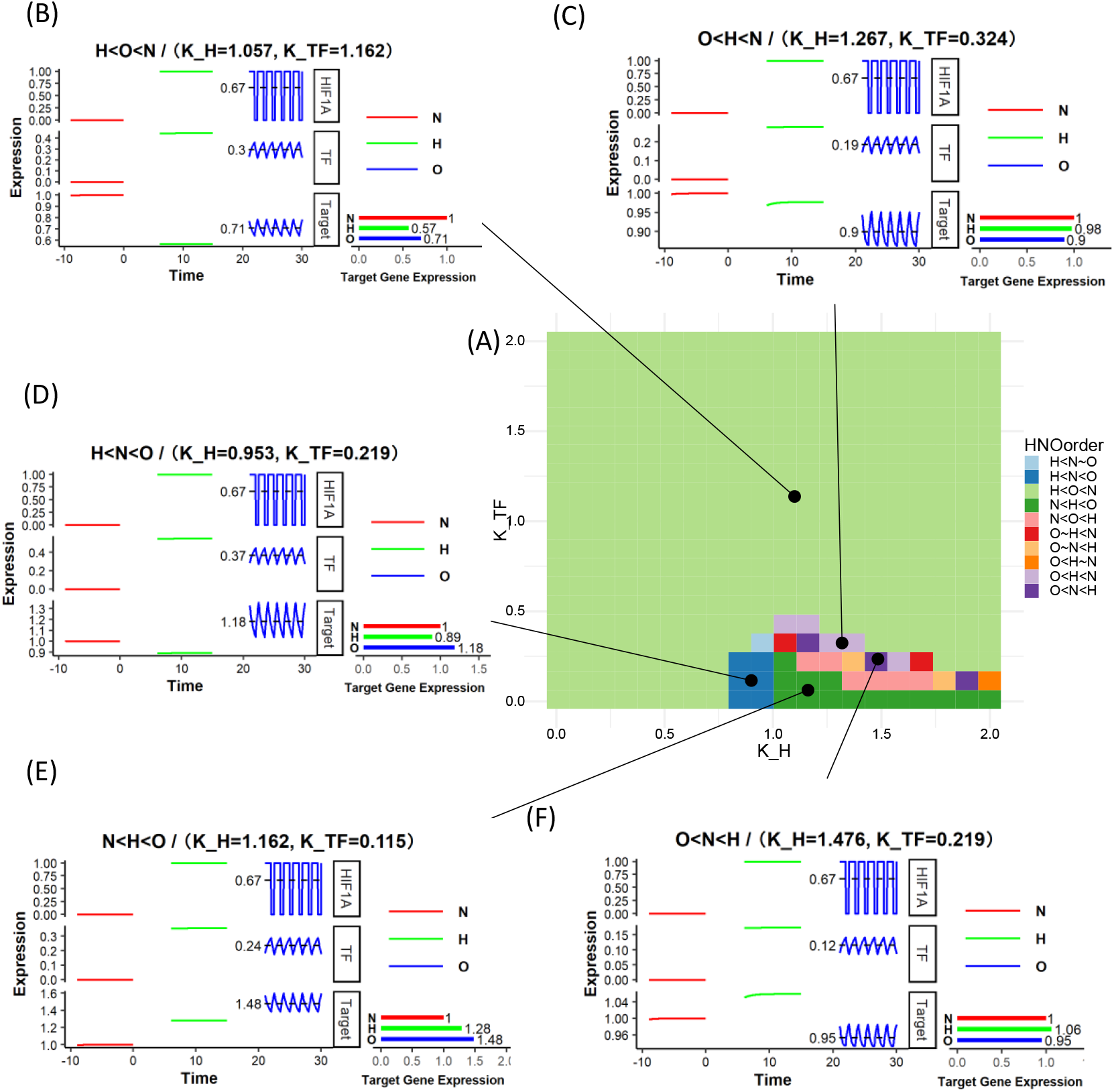
IFFLs with multiplicative effect of HIF-1 and HIF-1 activated TF on target gene expression show cycling/oscillatory hypoxia specific patterns. Phase diagram and example dynamics of IFFL Type 3 circuit with multiplicative inhibition (Equations 1 and 2). (A) Phase diagram showing the relative order of the target gene expression in Normoxia (N), Hypoxia (H), and Oscillatory Hypoxia (O). (B-F) Example dynamics of the driving HIF1A that is set at 1 for hypoxia, 0 for normoxia, and pulsating 1 hour of hypoxia and half an hour of normoxia for oscillatory hypoxia, the intermediate transcription factor, and the Target gene. All gene expression values are in arbitrary units. H, N, O order pattern given only in terms of the target gene expression.

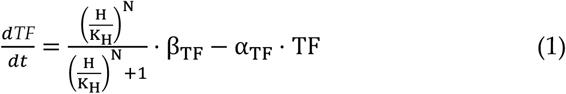

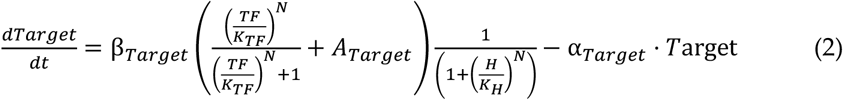

Where:

H: Concentration of HIF-1.

TF: Concentration of the transcription factor. Target: Concentration of the target gene product.

K_H_: Dissociation constant, indicates the concentration of HIF-1 at which the transcriptional response is at half-maximum.

*K*_*TF*_: Dissociation constant for TF at the target gene.

α_TF_: Degradation rate of TF.

β_TF_: Maximum production rate of TF.

α_*Target*_: Degradation rate of the target gene.

β_*Target*_ : Maximum production rate of the target gene influenced by TF.

*A*_*Target*_ : Additional activation of the target gene independent of TF.

N: Hill coefficient, describes the cooperativity of HIF-1 binding.

It can be observed that the silencing effect in Equation (2) is multiplicative to the HIF-1 driven enhancer. Physiologically, it pertains to the effect of HIF-1 and the induced TF to be transcriptionally synergistic, or cooperative^30^. Exploring IFFLs with the target gene with an independent inhibitory term, we obtain Equation (3).

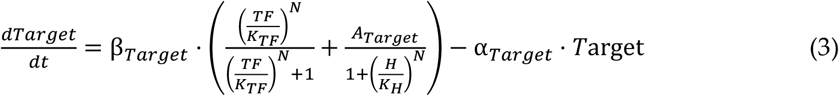

The additive silencing by the HIF-1 driven silencer is also modeled here, which gives rise to a very different phase diagram, wherein even larger parameter regimes show cycling hypoxia driven target gene expression.

### IFFL dynamics can generate gene expression responding specifically to cycling hypoxia

We sought to explore if a) our IFFL model can generate target gene expression that responds specifically to cycling hypoxia, and b) the parameter ranges that would be required for such emergent responses. We numerically solved the differential equations based models of IFFLs described above in R to simulate gene expression patterns in different hypoxic conditions and parameter ranges. We simulated the IFFL models both under multiplicative (**Figure 3**) and additive (**Figure 4**) inhibition, shown here for one of the IFFL sub-types (*type III*). We independently varied the HIF-1 dissociation constant (K_H_) and the intermediary transcription factor dissociation constant (*K*_*TF*_). At each parameter range, we classify the O specific responses according to the order of target gene expression under stable hypoxia (H), normoxia (N), and cycling/oscillatory hypoxia (O) (**Figure. 3A, 4A**).

**Figure 4.**
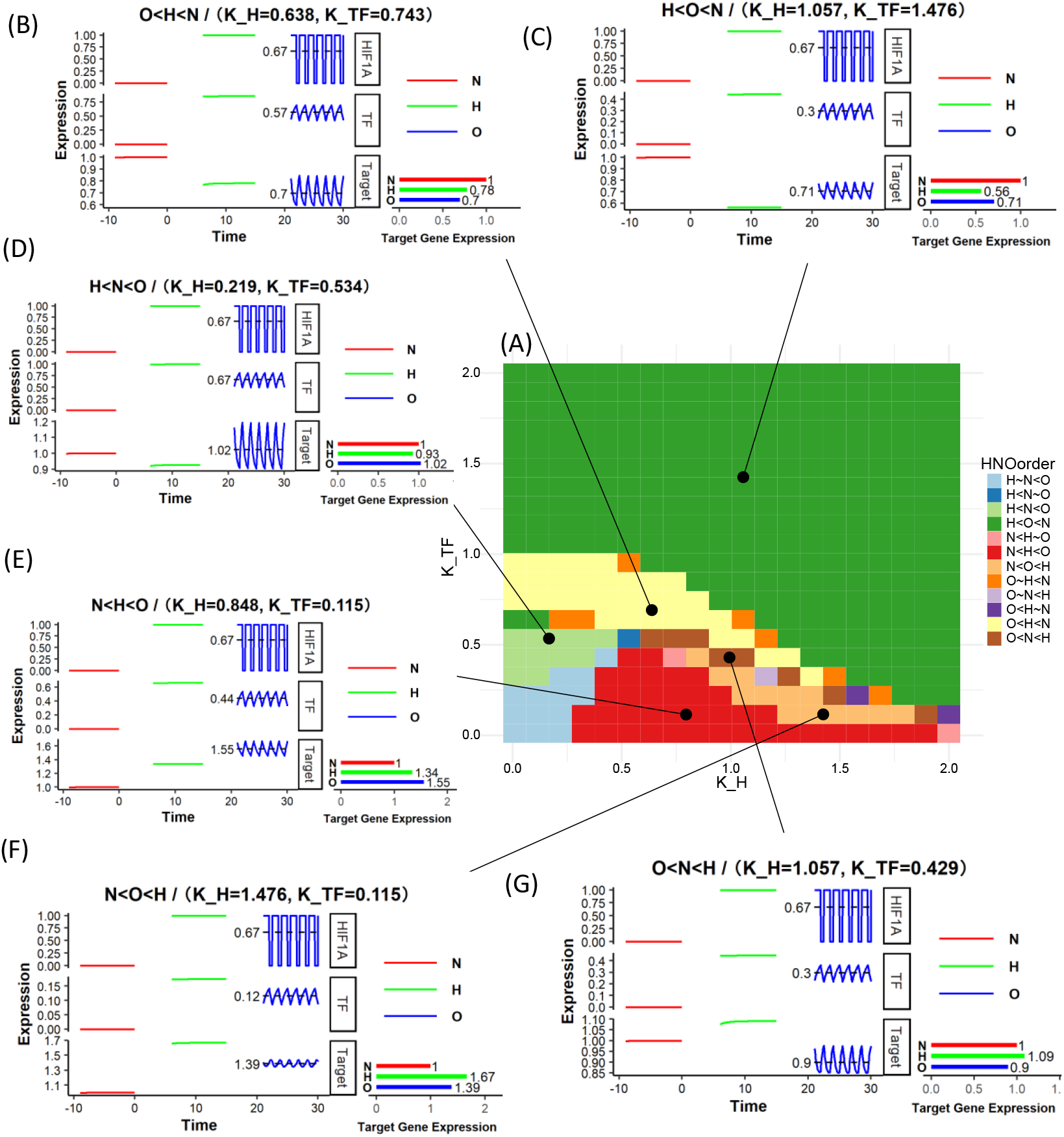
IFFLs with additive effect of HIF-1 and HIF-1 activated TF on target gene expression show cycling/oscillatory hypoxia specific patterns. Phase diagram and example dynamics of IFFL Type 3 circuit with additive inhibition (Equations 1 and 3). (A) Phase diagram showing the relative order of the target gene expression in Normoxia (N), Hypoxia (H), and Oscillatory Hypoxia (O). (B-G) Example dynamics of the driving HIF1A that is set at 1 for hypoxia, 0 for normoxia, and pulsating 1 hour of hypoxia and half an hour of normoxia for oscillatory hypoxia, the intermediate transcription factor, and the Target gene. All gene expression values are in arbitrary units. H, N, O order pattern given only in terms of the target gene expression.

HIF-1 is assumed to follow a cyclical abundance of 1 hour of stability followed by half an hour of destabilization, as shown in Figs. 3B-F and Figs. 4B-F. Specifically, to explore the relationship between target gene expression and the sensitivity of the system to hypoxia (*K*_*H*_) and TF activity (*K*_*TF*_), we simulated each of the four IFFL types, separately for multiplicative, and additive silencing by HIF-1. Phase diagrams identify distinct oscillatory/cycling hypoxia specific gene expression patterns. In the regimes where the inhibitory arm dominates, we observe, as in Fig. 3B and Fig. 4C, that the target gene expression is the lowest under hypoxia, intermediate in oscillatory hypoxia, and greatest under normoxia (H<O<N pattern). Similarly; when the activating arm dominates, as in Fig. 3E and Fig. 4F, we obtain the opposite, but expected pattern of N<O<H. However, for some parameter combinations wherein the effect of the activating and inhibiting arms are comparable but relatively staggered due to the nonlinearities of Equations 1-3, we observed the surprising behavior that the target gene expression under cycling hypoxia is either greater than under both hypoxia and normoxia (H<N<O) (**Figure 3D, 4D**), or (N<H<O) (**Figure 3E, 4E**), or lesser than both, as in Figs. 2C, 3B (O<H<N), Figs. 2F, 3G (O<N<H). For this IFFL circuit, within the same parameter range, the additive inhibitory model shows a larger regime of cycling hypoxia specific expression (**Figure 3**) than the multiplicative model (**Figure 4**).

We also computed similar phase diagrams for other IFFL circuits (**Figure 5**), showing substantial regimes of cycling hypoxia specific behavior (O<N<H, O<H<N, N<H<O, or H<N<O). Overall, our ODE based IFFL models demonstrate that cycling hypoxia specific gene expression patterns can emerge. Furthermore, IFFL models reflecting an additive effect of HIF-1 and HIF-1 driven TF, the parameter regimes for oscillation/cycling hypoxia specific gene expression was much larger than the multiplicative models.

**Figure 5.**
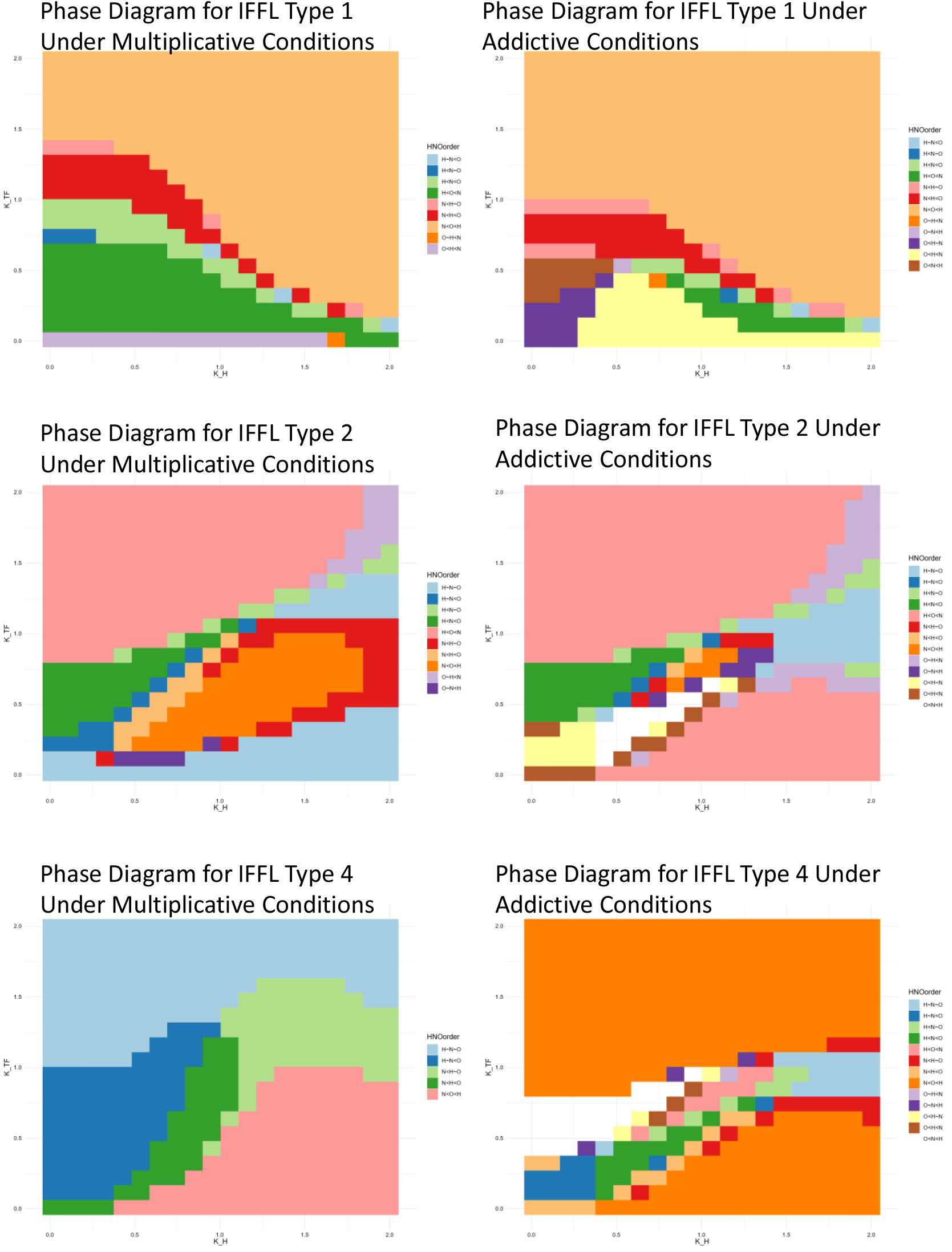
Both multiplicative and additive IFFLs show large parameter regimes with plausible oscillation specific gene expression patterns. Phase diagrams showing the H, N, O order for target gene expression under hypoxia, normoxia, and cycling/oscillatory hypoxia conditions for IFFL circuits of Type 1 (A), Type 2 (B), and Type 4 (C). The first column simulates under multiplicative inhibition (Equations 1 and 2), while the second column simulates the circuits under additive inhibition (Equations 1 and 3).

### HIF-1 recruits p53 and Notch1 in IFFL gene regulatory units to drive cycling hypoxia specific gene expression

We experimentally tested if IFFLs formed by HIF-1 are causal in oscillatory specific gene expression. Being a master transcriptional regulator^31^, HIF-1 transcriptionally activates a large number of genes, many of which themselves encode transcription factors. Based on a large number of previous reports, p53, and Notch1 were considered to be putative HIF-1 activated TFs, which could directly or indirectly influence gene expression of HIF-1 transcribed genes. Both p53, encoded by the TP53 gene, and Notch1 are important players in various cancers, and their involvement with HIF-is well established^32-34^. HIF-1 can reciprocally regulate p53, and also activates Notch1 by interacting with the intracellular domain of Notch (NICD)^33,35^.

To test if p53 and Notch1 can participate in IFFLs driven by HIF-1 and therefore regulate oscillatory/cycling hypoxia specific gene expression, we silenced TP53 and NOTCH1 gene in MDA-MB-231 cells, and sequenced RNA after the protocol adopted in normoxia (N), hypoxia (H), and cycling hypoxia (O). Filtering the genes that had significantly changed between N H O conditions in wild-type, and this pattern was disrupted in TP53 or NOTCH1 silenced conditions, we found several genes whose cycling hypoxia specific expression patterns were dependent on the participating TF. In case of TP53, we found all 4 classes of cycling/oscillatory hypoxia specific genes which were TP53 dependent on their unexpected expression patterns.

Several genes which showed patterns where cycling hypoxia had a directionally congruent, but a more extreme effect than stable hypoxia were found to be TP53 IFLL dependent (**Figure 5A-B**). Several of these TP53 dependent genes have potential role in cancer malignancies. These include SLC4A2, which encodes AE2, a bicarbonate-sodium exchanger which is upregulated in breast cancer contributing the metastasis^36^. We have previously also reported that oscillatory hypoxia results in increased expression of several ion channels and symporters^10,37^. Similarly, another key TP53 IFFL gene was ARHGAP39, a RHOA GTPase^38^ and ITFG2 which encodes Integrin alpha-2 implicated in breast cancer metastasis and poor survival outcomes^39^. Similarly, TBC1D2 which encodes a gene in the TBC domain family promotes E-cadherin degradation, and has a poor prognosis for ovarian cancer^40^. O<H<N pattern genes which were found to be TP53 dependent were mostly related to lipid metabolism, including FKBP7 a fatty acid binding protein, and HACD3 encoding for 3-hydroxyacyl-CoA dehydratase 3, as well as other pro-invasive genes THAP6 and LRRCC1, both associated with increased metastasis in various cancers (**Figure 5C**).

Interestingly, the genes which showed an opposite effect on expression in cycling hypoxia vs stable hypoxia were found to be more frequently TP53 dependent (**Figure 5D-E**). Among the genes which had H<N<O patterns in wild-type MDA-MB-231 and were found to be p53 dependent, most have been reported extensively for their role in TNBC (**Figure 5D**). Interestingly, many of these genes were associated with ER and Golgi based protein folding, or lipid metabolism, or those associated with autophagy, dysregulation of all critical in cancer progression. These included NFE2L1 which encodes the TF Nrf1, a key regulator of glucose and lipid metabolism^41,42^, MAP1LC3B encoding LC3B a key regulator of autophagy^42,43^, ERO1B encoding endoplasmic reticulum oxidoreductase 1 beta and strongly linked to TNBC malignancy and poor survival^44^, HSPA5 encoding heat shock protein family A member 5, and COG1 encoding a protein part of the conserved oligomeric golgi complex. In addition, EPAS1, which encodes HIF-2a and is linked to breast cancer prognosis was also found to be a gene which expresses in cycling hypoxia in a TP53 dependent manner^45^. Similarly, genes which increased in hypoxia, but decreased in stable hypoxia in wild-type cells but not in TP53 silenced cells included HPSE encoding heparinase, which is elevated in serum of breast cancer patients and has poor prognosis^46^, as well as NUDT9 which modulates HIF-1 activity (**Figure 5E)**^**47**^.

We also tested the hypothesis of HIF-1 driven IFFLs with another TF, Notch1, which is activated by HIF-1 itself and acts as a transcriptional regulator of many genes (**Figure 5F**). Using a similar methodology with TP53 KD cells in N, H, O, conditions, we identified a few genes that Notch1 may be regulating as part of IFFL with HIF-1. Again, we found genes that are increased in cycling hypoxia, although decreased in stable hypoxia, the pattern being Notch1 dependent, to be players in breast cancer progression These included NBPF12, CCDC9B, and GABRE, which encodes a subunit of GABA receptor. Overall, our experimental results not only identify, but demonstrate the causality of two key TFs which are HIF-1 regulated to participate in driving a gene expression pattern responsive specifically to oscillatory/cycling hypoxia, at times even opposite to stable hypoxia.

**Figure 6.**
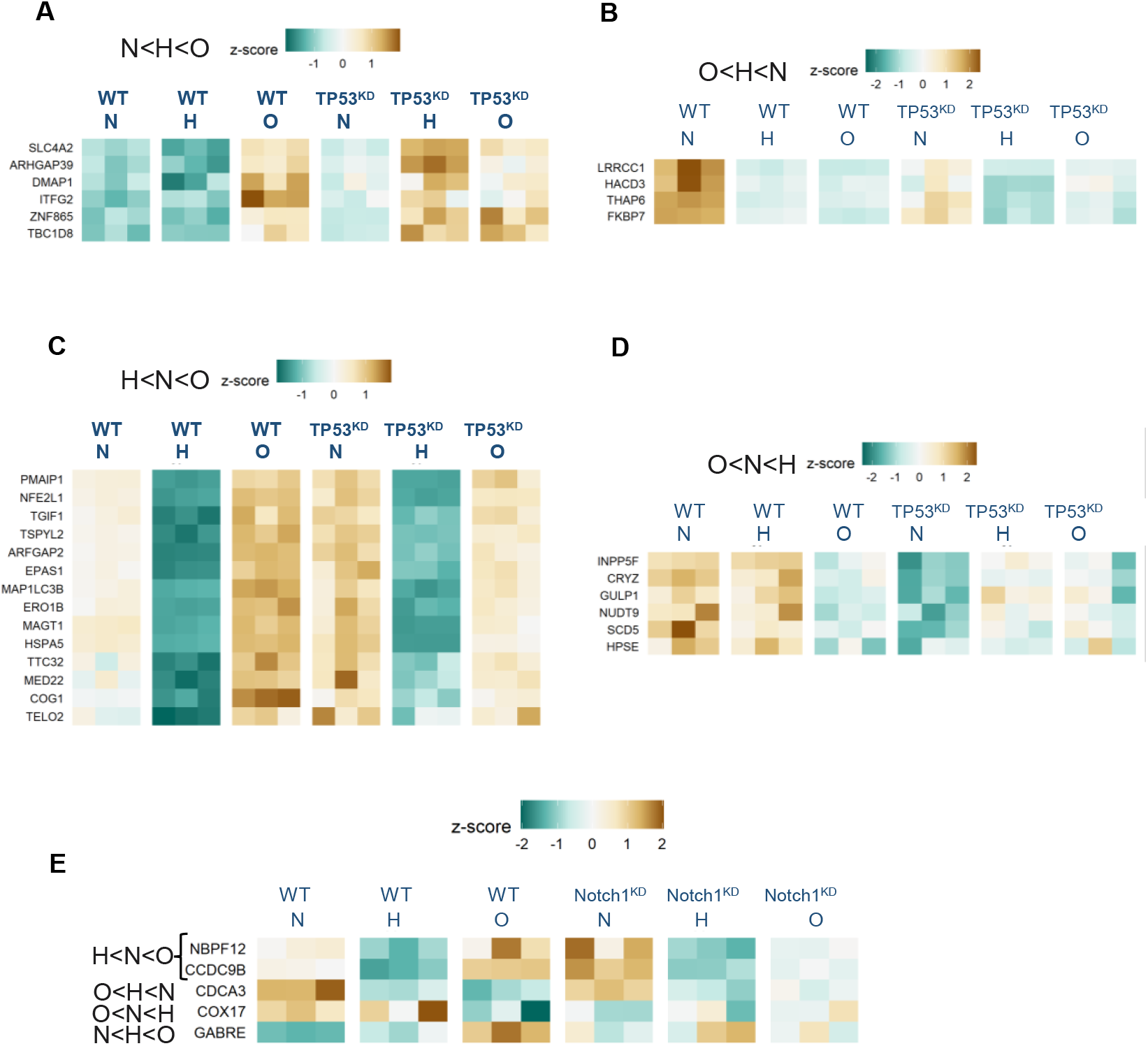
Target genes showing both oscillatory specific gene expression, and whose gene expression pattern can be explained by a gene circuit mediated by TP53 or Notch1. (A) N<H<O oscillatory specific pattern genes, whose oscillatory specific pattern is lost on TP53 knock down. (B) O<H<N oscillatory specific pattern genes, whose oscillatory specific pattern is lost on TP53 knock down. (C) H<N<O oscillatory specific pattern genes, whose oscillatory specific pattern is lost on TP53 knock down. (D) O>N>H oscillatory specific pattern genes, whose oscillatory specific pattern is lost on TP53 knock down. (E) Oscillatory specific pattern genes, whose oscillatory specific pattern is lost on Notch1 knock down.

## Discussion

Role of hypoxia is well established in nearly all aspects of tumor biology. Development of hypoxic cores in the growing tumors, wherein the vascular perfusion is unable to match the need for molecular oxygen, results in stabilization of HIF-1, a heterodimeric protein complex^2,6^. HIF-1 is a master transcriptional regulator, and its stabilization results in a remarkable change in cellular state, influencing many aspects of cellular metabolism, proliferation, differentiation, migration, and interaction with other cells. However, HIF-1 has also been shown to inhibit cell cycle progression, but cancers indeed continue to grow in hypoxia. We partly explained this conundrum by showing that fluctuations in HIF-1 stability, and therefore in downstream transcription, may provide the opportunity for cells to continue to divide even in hypoxia^14^. HIF-1 can exhibit oscillations in its transcriptional activity due to activation of chaperone mediated autophagy due to excess lactate in the hypoxic tumor milieu. In addition, various reports have shown that hypoxia itself is not stable in the tumor, and may cycle, particularly in the intermediate regions between the vascular and avascular regions^8,9^.

Here, we elucidate the plausible mechanism of specific gene expression pattern driven by hitherto less understood, or even appreciated aspect of tumor biology, cycling or oscillatory hypoxia^7^. Our RNAseq data on various cell types have revealed many genes which exhibit a cycling hypoxia specific gene expression pattern^10,14^. Ordinarily, cycling hypoxia signal would be averaged over many time periods to be in between the normoxic, and hypoxic range, and indeed, most genes exhibit this pattern. However, we found that for many genes, cycling/oscillatory hypoxia can increase the effect of hypoxia itself, even though the averaged O_2_ levels are lower when integrated over many cycles. Surprisingly, we also found that cycling/oscillatory hypoxia had an opposite effect on many downstream genes compared to stable hypoxia. These oscillation specific patterns are not possible to explain by simple gene regulation, suggesting more complex regulatory models.

We chose Incoherent Feed Forward Loop (IFFL) to model oscillatory input signals, and their effect on the target gene expression as plausible models for multiple reasons^20^. IFFLs are simple modules, consisting of just 3 factors, necessitating the inclusion of only one new species in addition to HIF-1 and the target gene. IFFLs are also a gene regulatory motif, indicating that they are evolutionarily selected owing to many selective advantages. Thirdly, because HIF-1 is a master transcriptional regulator, which regulates expression of many genes that themselves encode TFs, IFFLs are relatively easy to form with HIF-1. Finally, in certain parameter regimes, IFFLs can allow discrimination of variable and stable input signals^27^. We therefore mathematically modeled IFFLs to test the possibility of cycling/oscillation HIF-1 driven gene expression. All 4 IFFL models, integrating either the additive or synergistic/multiplicative of the direct effect of HIF-1 and HIF-1 induced TF on target gene, revealed large parameter regimes where oscillation specific gene expression was plausible. Interestingly, we found all specific subtypes of oscillation/cycling hypoxia driven gene expression patterns that we had observed in experimental data. The use of mathematical models to predict the behavior of these regulatory loops under varying hypoxic conditions opens new avenues for the development of precision therapies that can more effectively disrupt the adaptive mechanisms of cancer cells.

Finally, we identified two key TFs which could potentially participate in HIF-1 driven IFFLs to discriminate between variable and stable HIF-1 signals, with a profound effect on downstream gene expression. Both p53, and Notch1 are TFs critical in cancer progression, including breast cancer, and are known to be activated by HIF-1. Using gene perturbation of either TF encoding genes and subjected cells in normoxia, hypoxia, and cycling hypoxia conditions, we discovered several genes which are cycling/hypoxia specific in TP53, or Notch1 dependent manner. Remarkably, a large number of these genes, which putatively participate in IFFLs with HIF-1 and p53 (or Notch1) are pro-oncogenic genes, associated with poor prognosis in breast cancers, as well as other cancers.

More broadly, oscillatory signals are common across multiple scales of biology. Circadian rhythms, both systemic and the transcriptional clock regulators in the nucleus, endocrinal signals like the pulsating Gonadotropin releasing hormone production which drives the hyopthalamic-pituitary gonadal axis, as well as calcium waves within the cytoplasm are many examples of rhythmic, or cycling signals, critical for various physiological functions.

These cycling signals are processed in a specific way to elicit specific signaling, or gene expression responses. Their downstream processing itself could be complex, resulting in cyclical responses at a different frequency, e.g. hourly pulses of GhRH ultimately regulate the monthly ovulatory and menstrual cycles. Oscillations are certainly more common in signaling than are studied. Our approach presents an advancement in mechanistically understanding how oscillatory signals are processed downstream at the transcriptional level, presenting a methodological framework to identify the intermediaries which result in specific types of gene expression patterns. As many gene express in response to oscillatory signals, identification of other partners in the TF may provide new opportunities to therapeutically target transcription associated with various pathologies, including cancer. Finally, our work also pave a path to molecularly target cycling hypoxia specific gene expression by providing the putative transcriptional machinery underlying their expression.

## Methods

### Computational Modeling and Simulation of IFFL Circuits

To investigate the gene regulatory behavior of Incoherent Feed-Forward Loop (IFFL) circuits under varying oxygen conditions, we developed a mathematical model based on ordinary differential equations (ODEs) representing the interactions among HIF-1α (HIF1A), an intermediate transcription factor (TF), and a downstream target gene, as shown in Equations 1-3 and in the supplementary methods. Simulations were implemented using the deSolve package in R^48^.

For the generation of phase diagrams, we independently varied the dissociation constants for HIF1A and the intermediate TF (*K*_*H*_ and*K*_*TF*_) in a range from 0.01 to 2. This range was selected to ensure both mathematical stability and biological relevance, avoiding division by zero and preserving the numerical robustness of the simulations. Each parameter was discretized into 20 evenly spaced intervals. Unless otherwise noted, baseline parameter values, including degradation rates and maximum production rates of TF and the target gene (α_*TF*_, α_*Target*_,*β*_*TF*_, *β*_*Target*_), were set to 1.

To capture gene expression dynamics under normoxia (N), stable hypoxia (H), and cycling/oscillatory hypoxia (O), we selected representative points from the phase diagram for detailed time-series analysis. Simulations were conducted across a time interval from −15 to 30 arbitrary units with a resolution of 0.05. The intervals −15 to 0, 0 to 15, and 15 to 30 corresponded to normoxia, stable hypoxia, and cycling hypoxia conditions, respectively. To highlight the steady-state expression dynamics in each condition, we computed and visualized mean gene expression levels during the final 9 units of each interval: (−9, 0) for normoxia, (6, 15) for hypoxia, and (21, 30) for cycling hypoxia.

During the oscillatory phase, we modeled environmental cycling with alternating hypoxic (low) and normoxic (high) conditions. Each cycle consisted of 1 hour of normoxia followed by 0.5 hours of hypoxia, yielding a total cycle length of 1.5 hours, which approximates the experimental temporal dynamics applied in vitro.

### Transcriptional profiling of cancer cells

MDA-MB-231, a breast cancer cell line, was subjected to normoxia, stable hypoxia, and cycling hypoxia (1.5 hours of hypoxia and 0.5 hours of normoxia). We also obtained conducted CRISPR-mediated silencing of TP53 and NOTCH1 in MDA-MB-231 cells to obtain three different cell conditions: wild type (WT), TP53KD, and NOTCH1KD. This resulted in 3 cell conditions, with 3 oxygen treatments, and 3 technical replicates, with a total of 27 samples that were sequenced for mRNA profiling on the Illumina platform after standard mRNA library preparation. The obtained paired-end mRNA reads were aligned to the human genome using HISAT2^49^, and concordantly aligned counted against the standard gene models using featureCounts^50^. The sample x gene matrix was analyzed for differential expression using DESeq2^51^ on the R platform. Significance was defined as an adjusted p-value < 0.05. Genes exhibiting differential expressio^49^n across normoxic, hypoxic, and oscillatory conditions were identified and categorized by their N-H-O expression relationships.

To infer the regulatory role of TP53 or NOTCH1, we examined expression shifts in their respective knockout conditions. If a target gene Z showed a statistically significant increase in TP53KD or Notch1KD relative to WT, the repressive role of the corresponding transcription factor was inferred. Conversely, a decrease in expression indicated an activating relationship. Using this approach, we constructed putative HIF1A-TP53-target and HIF1A-Notch1-target regulatory loops based on congruent directional changes under HIF1A activation (assessed via expression in WT hypoxia versus normoxia).

## Acknowledgements

We are grateful to our funders: National Cancer Institute (1R37CA248161; PI: K), NICHD (K99HD105973; PI: YS), and AHA (24PRE1196464; PI: YL).

## Author Contribution

XQ wrote the code, performed bioinformatics analysis and experimental analysis; YL performed experiments on cycling hypoxia; EA, PNL helped in data interpretation; K conceptualized the project, provided funding, and wrote the manuscript; YS conceptualized and supervised the project, provided funding, and wrote the manuscript.

## Data Availability

The RNAseq data is available in NCBI Gene Expression Omnibus (GEO) database: Accession record: GSE292924. Code is deposited in GitHub: DOI: 10.5281/zenodo.15069799

## References

1 Pouyssegur, J., Dayan, F. & Mazure, N. M. Hypoxia signalling in cancer and approaches to enforce tumour regression. Nature 441, 437–443 (2006).

2 Wicks, E. E. & Semenza, G. L. Hypoxia-inducible factors: cancer progression and clinical translation. J Clin Invest 132 (2022).

3 Keith, B. & Simon, M. C. Hypoxia-inducible factors, stem cells, and cancer. Cell 129, 465–472 (2007).

4 Liu, S., Liu, Y., Qiu, X., Suhail, Y. & Kshitiz. Tissue-of-origin for cancers determines HIF-1 activation induced phenotypic heterogeneity. Mol Carcinog 63, 834–848 (2024).

5 Semenza, G. L. Hypoxia-inducible factor 1 (HIF-1) pathway. Sci STKE 2007, cm8 (2007).

6 Suhail, Y., Cain, M. P., Vanaja, K., Kurywchak, P. A., Levchenko, A., Kalluri, R. & Kshitiz. Systems Biology of Cancer Metastasis. Cell Syst 9, 109–127 (2019).

7 Michiels, C., Tellier, C. & Feron, O. Cycling hypoxia: A key feature of the tumor microenvironment. Biochim Biophys Acta 1866, 76–86 (2016).

8 Panek, R., Welsh, L., Baker, L. C. J., Schmidt, M. A., Wong, K. H., Riddell, A. M., Koh, D. M., Dunlop, A., McQuaid, D., d’Arcy, J. A., Bhide, S. A., Harrington, K. J., Nutting, C. M., Hopkinson, G., Richardson, C., Box, C., Eccles, S. A., Leach, M. O., Robinson, S. P. & Newbold, K. L. Noninvasive Imaging of Cycling Hypoxia in Head and Neck Cancer Using Intrinsic Susceptibility MRI. Clin Cancer Res 23, 4233–4241 (2017).

9 Matsumoto, S., Yasui, H., Mitchell, J. B. & Krishna, M. C. Imaging cycling tumor hypoxia. Cancer Res 70, 10019–10023 (2010).

10 Suhail, Y., Liu, Y., Du, W., Afzal, J., Qiu, X., Atiq, A., Vera-Licona, P., Agmon, E. & Kshitiz. Oscillatory hypoxia induced gene expression predicts low survival in human breast cancer patients. Mol Carcinog 63, 2305–2315 (2024).

11 Novin, A., Wali, K., Pant, A., Liu, S., Du, W., Liu, Y., Wang, L., Xu, M., Wang, B., Suhail, Y. & Kshitiz. Oscillatory Hypoxia Can Induce Senescence of Adipose-Derived Mesenchymal Stromal Cells Potentiating Invasive Transformation of Breast Epithelial Cells. Cancers (Basel) 16 (2024).

12 Du, W., Novin, A., Liu, Y., Afzal, J., Liu, S., Suhail, Y. & Kshitiz. Stable and Oscillatory Hypoxia Differentially Regulate Invasibility of Breast Cancer Associated Fibroblasts. bioRxiv (2024).

13 Korbecki, J., Siminska, D., Gassowska-Dobrowolska, M., Listos, J., Gutowska, I., Chlubek, D. & Baranowska-Bosiacka, I. Chronic and Cycling Hypoxia: Drivers of Cancer Chronic Inflammation through HIF-1 and NF-kappaB Activation: A Review of the Molecular Mechanisms. Int J Mol Sci 22 (2021).

14 Kshitiz Afzal, J., Suhail, Y., Chang, H., Hubbi, M. E., Hamidzadeh, A., Goyal, R., Liu, Y., Sun, P., Nicoli, S., Dang, C. V. & Levchenko, A. Lactate-dependent chaperone-mediated autophagy induces oscillatory HIF-1alpha activity promoting proliferation of hypoxic cells. Cell Syst 13, 1048–1064 e1047 (2022).

15 Hubbi, M. E., Hu, H., Kshitiz Ahmed, I., Levchenko, A. & Semenza, G. L. Chaperone-mediated autophagy targets hypoxia-inducible factor-1alpha (HIF-1alpha) for lysosomal degradation. J Biol Chem 288, 10703–10714 (2013).

16 Gardner, L. B., Li, Q., Park, M. S., Flanagan, W. M., Semenza, G. L. & Dang, C. V. Hypoxia inhibits G1/S transition through regulation of p27 expression. J Biol Chem 276, 7919–7926 (2001).

17 Koshiji, M., Kageyama, Y., Pete, E. A., Horikawa, I., Barrett, J. C. & Huang, L. E. HIF-1alpha induces cell cycle arrest by functionally counteracting Myc. EMBO J 23, 1949–1956 (2004).

18 Hubbi, M. E., Kshitiz Gilkes, D. M., Rey, S., Wong, C. C., Luo, W., Kim, D. H., Dang, C. V., Levchenko, A. & Semenza, G. L. A nontranscriptional role for HIF-1alpha as a direct inhibitor of DNA replication. Sci Signal 6, ra10 (2013).

19 Kshitiz, G. (figshare, 2025).

20 Mangan, S., Itzkovitz, S., Zaslaver, A. & Alon, U. The incoherent feed-forward loop accelerates the response-time of the gal system of Escherichia coli. J Mol Biol 356, 1073–1081 (2006).

21 Semsey, S. A mixed incoherent feed-forward loop allows conditional regulation of response dynamics. PLoS One 9, e91243 (2014).

22 Lamers-Lemmers, J. P., Hoofd, L. J. & Oeseburg, B. Oxygen diffusion coefficient and oxygen permeability of metmyoglobin solutions determined in a diffusion chamber using a non-steady state method. Adv Exp Med Biol 530, 509–517 (2003).

23 Saint-Andre, V. Computational biology approaches for mapping transcriptional regulatory networks. Comput Struct Biotechnol J 19, 4884–4895 (2021).

24 Kang, X., Hajek, B. & Hanzawa, Y. From graph topology to ODE models for gene regulatory networks. PLoS One 15, e0235070 (2020).

25 Seaton, D. D. ODE-Based Modeling of Complex Regulatory Circuits. Methods Mol Biol 1629, 317–330 (2017).

26 Semenza, G. L. A compendium of proteins that interact with HIF-1alpha. Exp Cell Res 356, 128–135 (2017).

27 Zhang, C., Tsoi, R., Wu, F. & You, L. Processing Oscillatory Signals by Incoherent Feedforward Loops. PLoS Comput Biol 12, e1005101 (2016).

28 Santos-Moreno, J., Tasiudi, E., Kusumawardhani, H., Stelling, J. & Schaerli, Y. Robustness and innovation in synthetic genotype networks. Nat Commun 14, 2454 (2023).

29 Rahi, S. J., Larsch, J., Pecani, K., Katsov, A. Y., Mansouri, N., Tsaneva-Atanasova, K., Sontag, E. D. & Cross, F. R. Oscillatory stimuli differentiate adapting circuit topologies. Nat Methods 14, 1010–1016 (2017).

30 Sanford, E. M., Emert, B. L., Cote, A. & Raj, A. Gene regulation gravitates toward either addition or multiplication when combining the effects of two signals. Elife 9 (2020).

31 Palazon, A., Goldrath, A. W., Nizet, V. & Johnson, R. S. HIF transcription factors, inflammation, and immunity. Immunity 41, 518–528 (2014).

32 Zhang, C., Liu, J., Wang, J., Zhang, T., Xu, D., Hu, W. & Feng, Z. The Interplay Between Tumor Suppressor p53 and Hypoxia Signaling Pathways in Cancer. Front Cell Dev Biol 9, 648808 (2021).

33 Wang, P., Guan, D., Zhang, X. P., Liu, F. & Wang, W. Modeling the regulation of p53 activation by HIF-1 upon hypoxia. FEBS Lett 593, 2596–2611 (2019).

34 Qiang, L., Wu, T., Zhang, H. W., Lu, N., Hu, R., Wang, Y. J., Zhao, L., Chen, F. H., Wang, X. T., You, Q. D. & Guo, Q. L. HIF-1alpha is critical for hypoxia-mediated maintenance of glioblastoma stem cells by activating Notch signaling pathway. Cell Death Differ 19, 284–294 (2012).

35 Guo, M., Niu, Y., Xie, M., Liu, X. & Li, X. Notch signaling, hypoxia, and cancer. Front Oncol 13, 1078768 (2023).

36 Khosrowabadi, E., Rivinoja, A., Risteli, M., Tuomisto, A., Salo, T., Makinen, M. J. & Kellokumpu, S. SLC4A2 anion exchanger promotes tumour cell malignancy via enhancing net acid efflux across golgi membranes. Cell Mol Life Sci 78, 6283–6304 (2021).

37 Liu, Y., Suhail, Y., Novin, A., Afzal, J., Pant, A. & Kshitiz. Lactate in breast cancer cells is associated with evasion of hypoxia-induced cell cycle arrest and adverse patient outcome. Hum Cell 37, 768–781 (2024).

38 Yao, L., Li, Y., Li, S., Wang, M., Cao, H., Xu, L. & Xu, Y. ARHGAP39 is a prognostic biomarker involved in immune infiltration in breast cancer. BMC Cancer 23, 440 (2023).

39 Cai, Y., Xu, G., Wu, F., Michelini, F., Chan, C., Qu, X., Selenica, P., Ladewig, E., Castel, P., Cheng, Y., Zhao, A., Jhaveri, K., Toska, E., Jimenez, M., Jacquet, A., Tran-Dien, A., Andre, F., Chandarlapaty, S., Reis-Filho, J. S., Razavi, P. & Scaltriti, M. Genomic Alterations in PIK3CA-Mutated Breast Cancer Result in mTORC1 Activation and Limit the Sensitivity to PI3Kalpha Inhibitors. Cancer Res 81, 2470–2480 (2021).

40 Tian, J., Liang, X., Wang, D., Tian, J., Liang, H., Lei, T., Yan, Z., Wu, D., Liu, X., Liu, S. & Yang, Y. TBC1D2 Promotes Ovarian Cancer Metastasis via Inducing E-Cadherin Degradation. Front Oncol 12, 766077 (2022).

41 Sekine, H. & Motohashi, H. Roles of CNC Transcription Factors NRF1 and NRF2 in Cancer. Cancers (Basel) 13 (2021).

42 Fernandez, L. P., Gomez de Cedron, M. & Ramirez de Molina, A. Alterations of Lipid Metabolism in Cancer: Implications in Prognosis and Treatment. Front Oncol 10, 577420 (2020).

43 Samdal, H., Sandmoe, M. A., Olsen, L. C., Jarallah, E. A. H., Hoiem, T. S., Schonberg, S. A. & Pettersen, C. H. H. Basal level of autophagy and MAP1LC3B-II as potential biomarkers for DHA-induced cytotoxicity in colorectal cancer cells. FEBS J 285, 2446–2467 (2018).

44 Varone, E., Decio, A., Chernorudskiy, A., Minoli, L., Brunelli, L., Ioli, F., Piotti, A., Pastorelli, R., Fratelli, M., Gobbi, M., Giavazzi, R. & Zito, E. The ER stress response mediator ERO1 triggers cancer metastasis by favoring the angiogenic switch in hypoxic conditions. Oncogene 40, 1721–1736 (2021).

45 Lu, X., Zhang, W., Zhang, J., Ren, D., Zhao, P. & Ying, Y. EPAS1, a hypoxia- and ferroptosis-related gene, promotes malignant behaviour of cervical cancer by ceRNA and super-enhancer. J Cell Mol Med 28, e18361 (2024).

46 Zahavi, T., Salmon-Divon, M., Salgado, R., Elkin, M., Hermano, E., Rubinstein, A. M., Francis, P. A., Di Leo, A., Viale, G., de Azambuja, E., Ameye, L., Sotiriou, C., Salmon, A., Kravchenko-Balasha, N. & Sonnenblick, A. Heparanase: a potential marker of worse prognosis in estrogen receptor-positive breast cancer. NPJ Breast Cancer 7, 67 (2021).

47 Yoon, B., Yang, E. G. & Kim, S. Y. The ADP-ribose reactive NUDIX hydrolase isoforms can modulate HIF-1alpha in cancer cells. Biochem Biophys Res Commun 504, 321–327 (2018).

48 Soetaert, K., Petzoldt, T. & Setzer, R. W. Solving Differential Equations in R: Package deSolve. Journal of Statistical Software 33, 1–25 (2010).

49 Kim, D., Paggi, J. M., Park, C., Bennett, C. & Salzberg, S. L. Graph-based genome alignment and genotyping with HISAT2 and HISAT-genotype. Nature Biotechnology 37, 907–915 (2019).

50 Liao, Y., Smyth, G. K. & Shi, W. featureCounts: an efficient general purpose program for assigning sequence reads to genomic features. Bioinformatics 30, 923–930 (2014).

51 Love, M. I., Huber, W. & Anders, S. Moderated estimation of fold change and dispersion for RNA-seq data with DESeq2. Genome Biol 15, 550 (2014).

